# An optic ray theory for durotactic axon guidance

**DOI:** 10.1101/2020.12.16.423083

**Authors:** Hadrien Oliveri, Kristian Franze, Alain Goriely

## Abstract

During the development of the nervous system, neurons extend bundles of axons that grow and meet other neurons to form the neuronal network. Robust guidance mechanisms are needed for these bundles to migrate and reach their functional target. Directional information depends on external cues such as chemical or mechanical gradients. Unlike chemotaxis that has been extensively studied, the role and mechanism of durotaxis, the directed response to variations in substrate rigidity, remain unclear. We model bundle migration and guidance by rigidity gradients by using the theory of morphoelastic rods. We show that at a rigidity interface, the motion of axon bundles follows a simple behavior analogous to optic ray theory and obeys Snell’s law for refraction and reflection. We use this powerful analogy to demonstrate that axons can be guided by the equivalent of optical lenses and fibers created by regions of different stiffnesses.

---

The establishment of the neural network is an event of paramount importance in brain development. During this process, neurons extend slender processes called *axons* must grow, possibly to great length, along precise pathways in search of their functional target. To establish connections across different regions, axons group themselves into bundles and migrate together until they reach their target where they split [1]. The role of chemical cues in axon pathfinding is well established, and different important chemo-attractants and repellents have been identified [2–4]. Chemical gradients are perceived by the sensory tip of each axon and integrated as directional cues for pathfinding [4]. This mechanism is the basis for many theoretical models of axon navigation [5–7, 9–14]. In contrast, the role of mechanical cues in guidance has received little attention [15]. In particular, *durotaxis*, the directed motion or growth of cells based on variations in the stiffness of their extracellular matrix has only been discovered recently [16, 17], but is believed to be of critical importance for guidance [18–20]. Understanding *durotactic guidance* requires detailed measurement of heterogeneous tissue stiffness properties [18, 20–24], as well as a mechanistic understanding of axon locomotion and mechanics [25].

Mechanically, axon migration and elongation are mediated by the *growth cone*, the actomyosin-rich distal structure of the axon. The growth cone uses focal adhesion and active contractile forces to pull on the extracellular substrate [15]. This contraction generates tension in the trailing axon shaft, which responds to stress by yielding anelastically [26]. Axons in a bundle bind to one another via membrane-membrane adhesion forces [27]. Depending on its individual position in the fascicle, each growth cone perceives a different signal, and thereby exerts a slightly different traction. These forces add up to produce collectively a global *wrench* (the simultaneous application of a force and a torque) acting at the tip of the bundle structure (Fig. 1, inset). The ability of growth cones to grip onto their medium and produce traction is enhanced on stiffer substrates; therefore, differences in substrate rigidity on the bundle-width length scale, result in emergent durotaxis through bundle deflection toward *softer* medium [18]. Following this principle, a variation in stiffness induces locally a wrench at the tip of the axon bundle. The question is then to characterize the global motion of the bundle, and to understand how substrate rigidity can be used as a guidance mechanism.

**FIG. 1.**
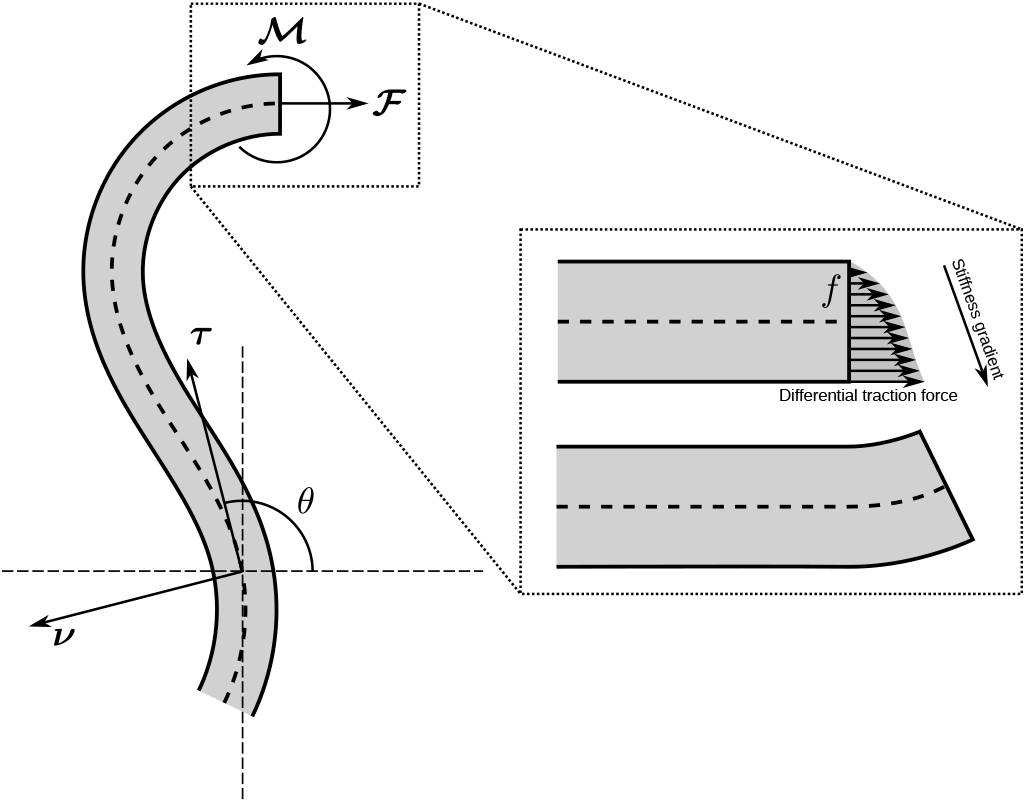
An axon bundle is modeled as a rod subject to a wrench 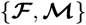. Inset: The wrench is induced by the forces generated by individual axons. The unit tangent ***τ*** = ∂***r***/∂*s* = (cos *θ,* sin *θ*) and normal ***ν*** = (−sin *θ,* cos *θ*) vectors are defined with respect to the central axis through the angle *θ*.

## Model

We are interested in the motion of the entire bundle away from fasciculation and defasciculation events [28]. In this regime, the bundle behaves mechanically as a single rod (Fig. 1). Therefore, we model the axon bundle as a single unshearable planar *morphoelastic rod* [29, 30] subject to a wrench given by a torque 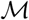 and a resultant 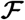 applied at the tip. We assume that each growth cone mostly pulls along the current axis of the bundle (Fig. 1, inset). This is sufficient to produce durotactic turning, which emerges as a collective effect, rather than from autonomous reorientation of each single axon. Therefore, bundle reorientation is caused by the torque 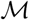, whereas the resultant 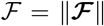 only affects the growth speed. Geometrically, the bundle is characterized at time *t* by a smooth curve oriented from the base to the tip: ***r*** (*s*) = (*x* (*s*) *, y* (*s*)), where *s* is the current arc length (*SI*, Section I).

We account for axon-substrate adhesion through friction forces that work against the bundle shaft motion [31]. Consequently, the tension applied by the growth cone dissipates into the substrate, so that only a finite distal section of the tip effectively grows (*SI*, Section I E). Therefore, in the limit of strong adhesion considered here, bundle migration is a tip growth process and we can model the bundle path as a tip-growing curve fully parameterized by its arc length *s*, and curvature, *κ* = *∂θ/∂s*, given by

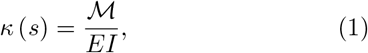

where *E* and *I* are the bundle’s Young’s modulus and second moment of area. The tip torque 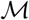 depends on the *local* substrate rigidity *C* = *C* (*X, Y*) (with the dimension of pressure). An infinitesimal surface element d*A* in the tip cross section is subject to the longitudinal force 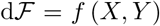, due to the traction density *f* (*X, Y*). For typical tissue stiffnesses measured *in vivo* (*C* ≃ 0.1 kPa) [18], we can neglect frictional slippage, that result in loss of grip on stiffer substrates (*C* > 1 kPa) [25]. Assuming small deformations, the traction is linearly coupled to the rigidity as *f* = *ϑC*, where *ϑ* is a dimensionless constant. Therefore, the integration of the force density over the tip cross section provides the total wrench applied to the rod:

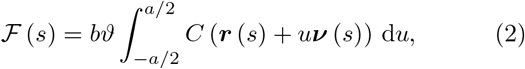

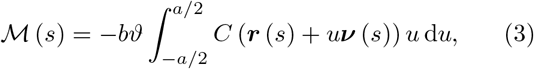

where *b* is the out-of-plane thickness of the bundle; *a* = 1 its in-plane contact width, taken as a reference length; and *u* ∈ [−*a/*2, *a/*2] is a cross-sectional coordinate in the direction of the unit normal vector ***ν*** (Fig. 1). For a graded smooth stiffness field, this wrench may be approximated to leading order by (*SI*, Section II A)

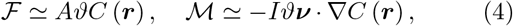

where we see clearly the effect of a stiffness gradient on the torque exerted on the bundle tip.

## Laws of refraction

We first consider the canonical problem of an axon bundle entering a straight interface between two media, by taking *C*(*X, Y*) = *C*_1_ if *X* < 0, and *C*(*X, Y*) = *C*_2_ otherwise. Then, Eqs. (2) and (3) simplify to

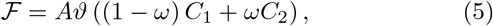

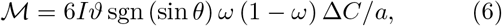

where *A* = *ab*; *I* = *a*^3^*b/*12; Δ*C* = *C*_2_ − *C*_1_; and the indicator function *ω* = *ω* (*x, θ*) ∈ [0, 1] codes the proportion of the tip width that is inside the *C*_2_ zone (*SI*, Section II B). A torque is produced when the bundle tip is in contact with the interface, i.e. when *ω* (1 − *ω*) > 0; outside this region of influence, there is no torque and the motion follows a straight line (Fig. 2A). The angular deflection of the bundle due to the interaction between the bundle and the interface, is governed by

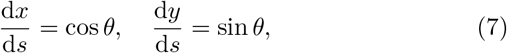

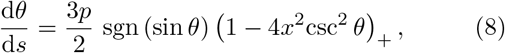

where *p* = *ϑ*Δ*C/E*, the *durotactic number*, is dimensionless; and (*x*)_+_ ≔ max (0, *x*) is the ramp function. A first integral of this system can be obtained exactly (*SI*, Section III), and its level sets are illustrated in the *x*-*θ* phase plane of Fig. 2B. By integrating the trajectory while the bundle tip is in contact with the interface (red and green zones of Fig. 2B), we find that the deflection between the two angles of incidence *θ*_1_ and *θ*_2_ (Fig. 2A, inset) obeys a Snell-type law [32] of the form

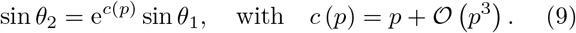

**FIG. 2.**
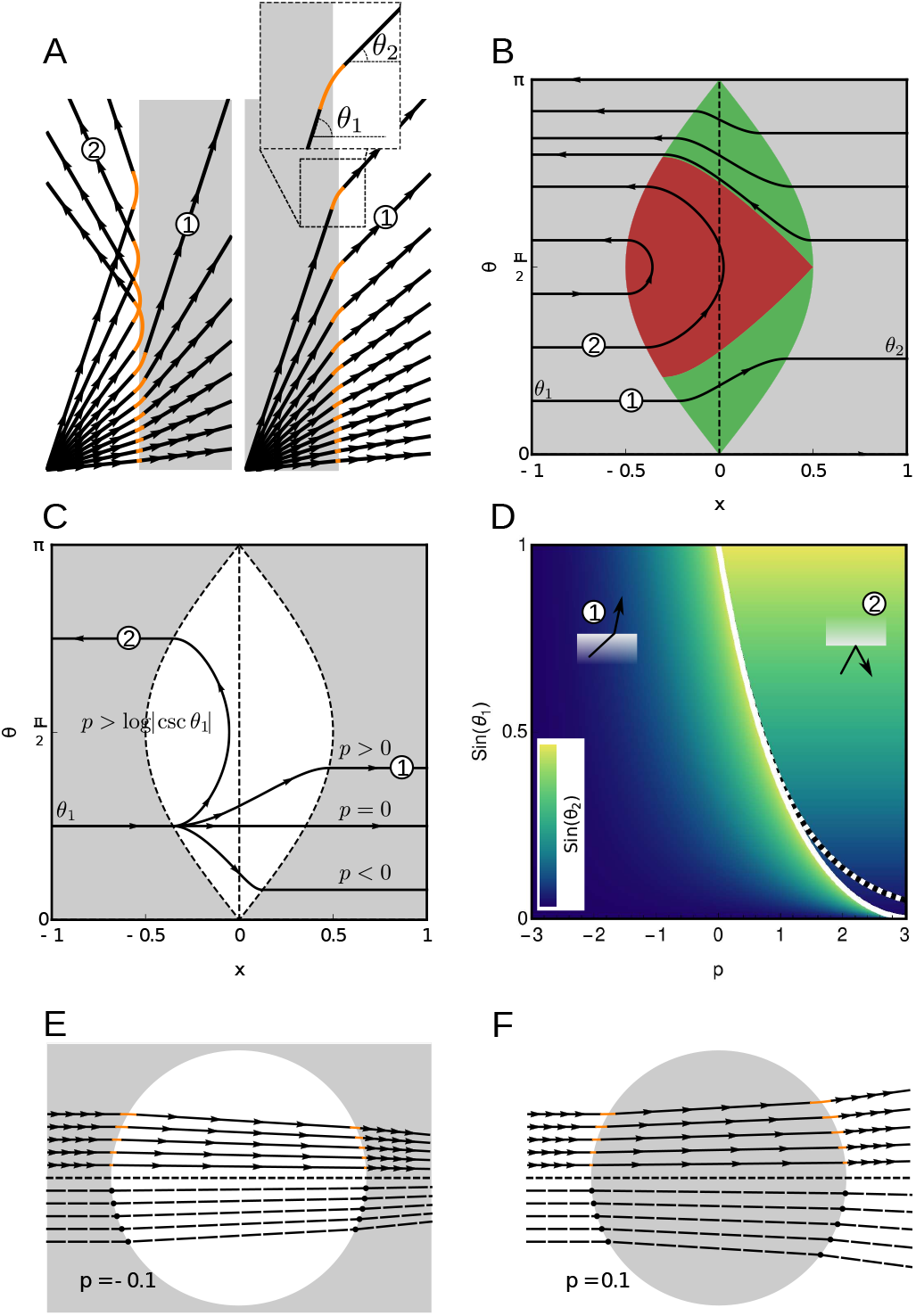
Durotactic refraction (labeled by 1) and reflection (2). (A) Trajectories of axon bundles through a hard interface (*p* = 0.25). Orange indicates the portion of the trajectory where the torque is non-zero. Gray and white zones indicate stiff and soft domains. (B) Phase portrait of System (7–8) (*p* = 0.5). Gray zone corresponds to the *θ*-nullcline set, i.e. *ω* = 0 or 1. Red and green domains depict the sets of reflection and refraction paths. (C) Trajectories for different *p* (with *θ*_1_ = *π/*4). When *p* exceeds the critical value ≃ log |csc *θ*_1_|, the system bifurcates from a refraction to a reflection regime. (D) Parameter space sin *θ*_1_ vs. *p*. Heat map: sin *θ*_2_. White and dashed lines indicate the critical line exp (−*c* (*p*)) and its approximation exp (−*p*). (E, F) Convergent and divergent lensing effect produced by a soft or stiff circular obstacle (diameter = 10, *p* = ±0.1). Solid lines are obtained by integrating the equations of motion and dashed lines are the geometric rays from the Snell approximation.

Therefore, for |*p*| ≪ 1, we obtain the familiar approximation

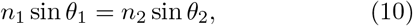

where *n*_*i*_ = exp (−*ϑC*_*i*_/*E*), is the refractive index of medium *i*. Bundle reflection (*θ*_2_ = *π* − *θ*_1_) occurs when a bundle migrates from a soft to a stiff medium (*n*_2_ < *n*_1_) with an incidence angle |*θ*_1_| larger than the critical angle *θ*\* = arcsin (*n*_2_/*n*_1_) (Fig. 2C and D).

The behavior of bundles at a stiffness jump matches *precisely* the fundamental rules governing light ray deflection at a refractive jump [32]. Since this property only depends on the local geometry of an interface, the result is naturally extended to any interface that does not vary significantly on the bundle-width length scale, as shown next.

## Durotatic lenses

The striking formal analogy between optic ray theory and axon motion automatically implies that all results of geometric optics based on Snell’s law can be formally applied to our problem, notwithstanding the fundamental differences between light propagation and axon growth. For instance, we can simulate the lensing effects created by a circular interface, as illustrated in Fig. 2E and F. Here, a soft obstacle acts as a convergent lens (Fig. 2E), whereas a stiff obstacle acts as a divergent lens (Fig. 2F). To validate our approximation, we both show trajectories obtained by direct integration of the mechanical problem (Eqs. (7) and (8)), and by a geometric construction based on the durotactic Snell law (Eq. (10)). These idealized examples show that durotactic effects and their associated Snell’s law can be used to guide bundle trajectories by controlling the spread or focus of axon bundles.

## Duroducts

The theory proposed so far is idealized as it neglects the stochastic behavior of axons subject to noise and imperfections in their ability to sense and respond to tissue stiffness. To establish the neural network, axon guidance must be sufficiently robust to ensure that a bundle reaches its target. A similar problem arises in light-based communication technologies, where light needs to be carried over large distances with minimal loss. An elegant technical solution to this requirement is provided by *optical fibers*, slender refractive tubes in which light is guided by internal reflections [33]. We adapt this idea to durotaxis by considering the motion of a growing bundle in a *duroduct* : a soft corridor of characteristic width 2*R* between two stiff regions (Fig. 3). Here we use a graded smooth stiffness field with a wrench given by Eq. (4). To account for the random component of axon movement, we add a Gaussian noise to the curvature *κ* (*s*) (Eq. (1)).

**FIG. 3.**
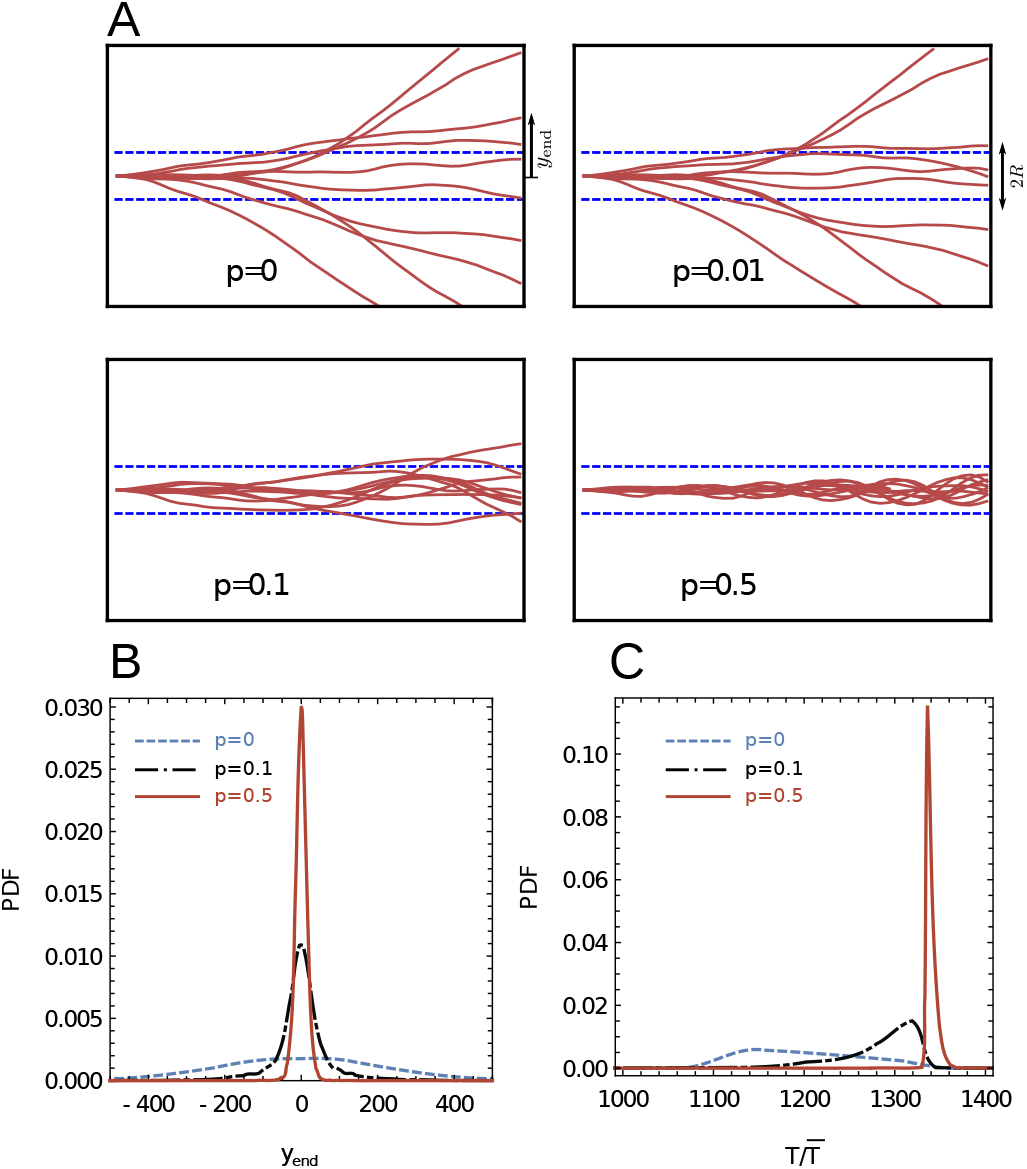
Noised migration of axon bundles in a *duroduct*. (A) Plot of *N* = 10 representative trajectories obtained for various values of *p*. Dashed blue lines show the characteristic width 2*R* of the soft corridor (*R* = 50 in this example). (B, C) Estimated probability density function (PDF) of (B) the final coordinate *y*_end_ and (C) normalized travel time 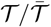 (*SI*, Section IV) for *p* = 0, 0.1 and 0.5 (*N* = 10^4^ realizations).

We consider the motion of a bundle from left to right, starting at *x* (0) = *y* (0) = *θ* (0) = 0 until it reaches *x* = 1000 at which point, its coordinate *y* = *y*_end_ is recorded (Fig. 3A). Multiple realizations of this motion provide a statistical distribution for *y*_end_ (Fig. 3B).

In the absence of a duroduct (Fig. 3A, *p* = 0), noise will cause most of the bundles to diverge away from the central axis. Consequently, the probability of an axon to stay within the corridor (|*y*_end_| ≤ *R*), will decrease with the travel distance. Increased durotactic number *p >* 0 mitigates the effect of noise by decreasing the probability of bundle loss, and sharpens the distribution around the axis (Fig. 3B). Remarkably, for larger *p* (e.g. *p* = 0.5), the duroduct effectively acts like an optical fiber, since the majority of the bundles are trapped and bounce against the soft boundaries of the confinement zone (see also *Supp. Movie* S1). This behavior indicates that a duroduct has the potential to confer spatial robustness to long-distance axon migration. We also show that the probability density of travel times also sees its variability decrease as *p* increases (Fig. 3C).

## Xenopus optic tract

As an application of these ideas, we examine the case of durotactic guidance of xenopus *retinal ganglion cell* (RGC) axons, located in the retina [18, 20]. RGC axons leave the retina in the optic nerve, which crosses the midline at the optic chiasm. Axons then grow along the contralateral brain surface towards the optic tectum, where they terminate. A gradient in brain tissue rigidity is observed, which correlates strongly with the stereotypical caudal turn undergone by RGC axons in the mid-diencephalon on their way to the tectum [18]. We test the possible role of durotaxis in caudal turn using our optic model of optic tract on a brain stiffness map obtained from atomic force measurements [20] (*SI*, Section V A). To simulate growth and guidance, we initiate *N* = 10^4^ axon rays, starting from the entry point of the domain (bottom right corner in Fig. 4A), with a variability on initial position and angle (dashed orange hull in Fig. 4A), and noise on curvature. We define the target zone as the quadrant represented by a dashed line in Fig. 4A, and for each value of *p* tested, we record the success ratio *N*_*p*_/*N*, with *N*_*p*_ the number of tracts that reach the target (Fig. 4B).

**FIG. 4.**
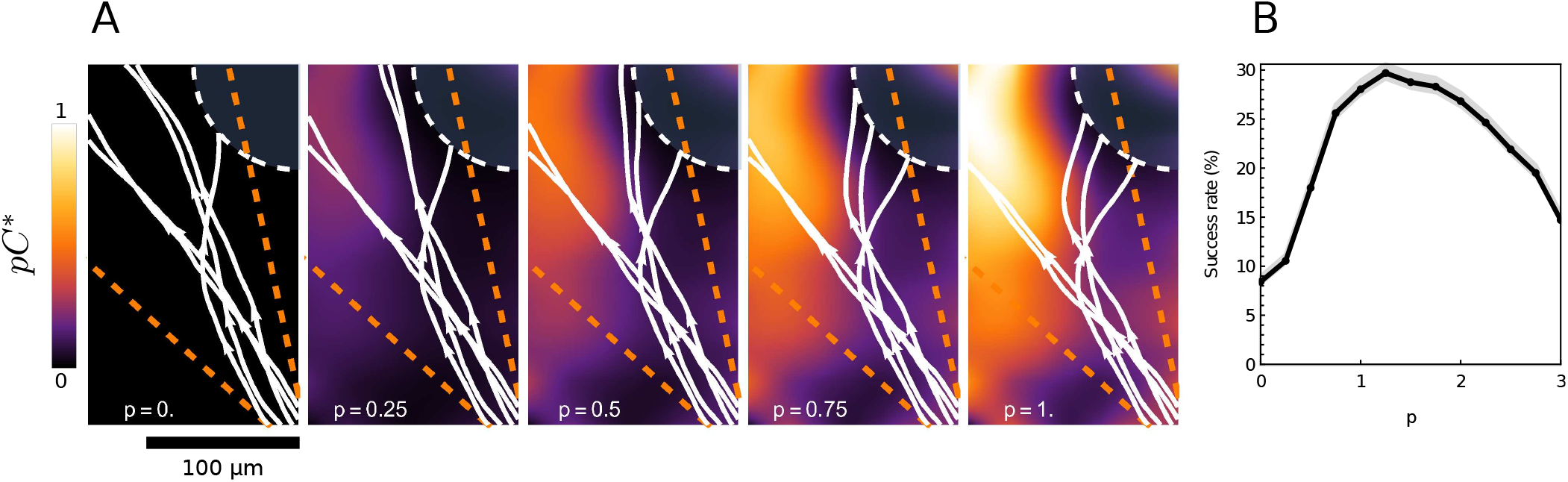
Simulation of the xenopus optic tract. (A) Plot of *N* = 5 representative trajectories (white solid lines) for different *p*. Target is represented by the quarter circle (white dashed line). Orange dashed lines define the set of ballistic paths, obtained in the absence of noise and durotaxis. Heat map corresponds to product *pC*\* where *C*\* is the medium stiffness normalized to [0, 1] (*SI*, Section V A). (B) Success rate *N*_*p*_/*N* vs. *p* (*N* = 10^4^ realizations). Gray streak: 95% confidence intervals.

Without durotaxis (*p* = 0), more than 90% of the unguided tracts miss the target. However, as *p* increases, a clear increase in success rate is observed, with a peak at *p* ≃ 1. Here, many tracts undergo a deflection that resembles the turn seen *in vivo*, confirming that durotaxis has the potential to contribute to caudal turn. From dimensional analysis (*SI*, Section V B), we expect the durotactic number *p* in this experiment to be on the order of 0.01–1, and may be sufficient for the bundle to reach its target *in silico* albeit not with the sharp turn observed *in vivo*. Therefore, durotaxis, on its own, is not fully sufficient to account for the *in vivo* observations, and other processes like chemorepulsion [34–36], and steric hindrance due to higher cell body densities in the mid-diencephalon [18] should be combined to obtain a full picture of guidance.

## Discussion

Axon guidance is a complex mechanism that relies on many physical and chemical cues. Clearly, durotaxis by itself may not be sufficient to establish a functional network as it lacks the specificity needed to find a precise cellular target. Yet, the prepatterning of tissues with different stiffnesses in the nervous system may provide a universal mechanism to aid guidance through durotaxis without the need to maintain chemical gradients over large distance during development. Furthermore, we showed that the motion of axon bundles due to variation in stiffness follows a simple refraction law, and that proper patterning can enhance the precision of guidance by narrowing the distribution of axon endpoints around a target. In addition, this narrowing of the distribution in space implies a narrowing of the distribution in the times at which a target is reached. Robustness in timing is a fundamental issue during neurodevelopment when a number of developmental events must happen in a precise order and in synchronicity [37].

We see the analogy with the geometric light rays as potentially powerful. Indeed, we know from the theory of optics that ingenious devices such as lenses, mirrors, optical guides, collimators, binoculars, periscopes, telescopes, and microscopes can be built to control the path of light rays and collect information. Whether similar instruments can be found in nature or devised *in vitro* remains to be established. Yet, the possibility that our knowledge of optics can be ported to the motion of axons opens new avenues of research.

The support for A.G. by the *Engineering and Physical Sciences Research Council* of Great Britain under research grant EP/R020205/1 is gratefully acknowledged. K.F. acknowledges funding from the *European Research Council* (Consolidator Award 772426), and the *Alexander von Humboldt Foundation* for his Alexander von Humboldt Professorship. The authors also thank Ryan Greenhalgh from the University of Cambridge, for insightful discussions and for providing access to the data.

