## Supplementary Information for "An optic ray theory for durotactic axon guidance"

Hadrien Oliveri[1], Kristian Franze[2, 3] and Alain Goriely[1*]
[1] *Mathematical Institute, University of Oxford, Oxford, United Kingdom*
[2] *Department of Physiology, Development and Neuroscience, University of Cambridge, Cambridge, United Kingdom* and
[3] *Friedrich-Alexander-Universität Erlangen-Nürnberg, Max-Planck-Zentrum für Physik und Medizin, Erlangen, Germany*
(Dated: December 16, 2020)

### I. ROD MODEL

To model the growth of an axon bundle, we use the theory of morphoelastic rods that generalizes the classic Kirchhoff-Cosserat theory of filaments to a theory of growth and remodeling [1, 2]. We model the axon tract as a planar, unshearable, twistless, growing rod.

#### A. Geometry

In two dimensions, a twistless rod is completely specified by a smooth parametric curve $S_0 \in [0,1] \to \boldsymbol{r}(S_0) = (x(S_0), y(S_0)) \in \mathbb{R}^2$. The arc length $s(S_0, t)$ is oriented from the base ($s(0,t) = 0$) to the tip ($s(1,t) = \ell(t)$). The rod geometry can be described from the curvature $\kappa(s)$, by integration of the Frenet-Serret equations:

$$\frac{\partial x}{\partial s} = \cos\theta, \quad \frac{\partial y}{\partial s} = \sin\theta, \quad \frac{\partial \theta}{\partial s} = \kappa, \tag{S1}$$

where $\theta$ is the angle between the tangent vector $\boldsymbol{\tau} = \partial \boldsymbol{r}/\partial s = (\cos\theta, \sin\theta)$ and the $x$-axis. The normal vector is $\boldsymbol{\nu} = (-\sin\theta, \cos\theta)$, and $\lambda := \partial s/\partial S_0 > 0$ is the total stretch.

#### B. Balance of momentum

Mechanically, the rod is subject to internal and external forces. The internal forces at a given point $S_0$ are given by the resultant $\boldsymbol{n} = F\boldsymbol{e}_X + G\boldsymbol{e}_Y$ and the bending moment $\boldsymbol{m} = m\boldsymbol{e}_Z$ applied to the cross section $S_0$ by the distal section of the rod. In addition, we introduce external frictional forces due to adhesion with the substrate, characterized by a friction coefficient $\zeta$ [3]. Neglecting inertial effects, the balance of linear and angular momenta reads [2]:

$$\frac{\partial F}{\partial s} = \zeta \frac{\partial x}{\partial t}, \quad \frac{\partial G}{\partial s} = \zeta \frac{\partial y}{\partial t}, \tag{S2}$$

$$\frac{\partial m}{\partial s} = F\sin\theta - G\cos\theta. \tag{S3}$$

#### C. Constitutive laws

To close the problem, we need constitutive assumptions on the material. We model the elastic and anelastic properties of the bundle through the standard multiplicative decomposition [1]

$$\frac{\partial s}{\partial S_0} = \frac{\partial s}{\partial S}\frac{\partial S}{\partial S_0} \quad \Leftrightarrow \quad \lambda = \alpha\gamma, \tag{S4}$$

where $\alpha$ is the elastic stretch; $\gamma$ is the growth ratio; and parameter $S$ depicts positions in the intermediate (grown) configuration. We postulate that, constitutively, only $\alpha$ generates extensional stress, while, by contrast, $\gamma$ characterizes the grown reference configuration.

To model elasticity, we use a standard energy-density function (energy per unit intermediate length) $W(\alpha, \kappa) := EA(\alpha - 1)^2/2 + EI\kappa^2/2$, which corresponds to the energy of a naturally straight linear rod with Young's modulus $E$; cross-sectional area $A$; and second moment of area $I$. The constitutive equations for the extensional and bending responses are:

$$n = \frac{\partial W}{\partial \alpha} = EA(\alpha - 1) \tag{S5}$$

where $n := F\cos\theta + G\sin\theta$ is the longitudinal tension; and

$$m = \frac{\partial W}{\partial \kappa} = EI\kappa. \tag{S6}$$

Growth of the shaft is accounted for by increasing the growth multiplier $\gamma$. Several more or less detailed stress-based or strain-based laws have been proposed to model single axon growth [2–9]. Here, we use a minimal stress-based exponential growth law [3, 7]:

$$\frac{1}{\gamma}\frac{\partial \gamma}{\partial t} = \frac{n}{A\eta}. \tag{S7}$$

#### D. Boundary conditions

The boundary conditions at $S_0 = 1$ account for the wrench produced by the growth cones and applied to the trailing bundle

$$\boldsymbol{n}\big|_{1,t} = \boldsymbol{\mathcal{F}}(t), \quad m\big|_{1,t} = \mathcal{M}(t). \tag{S8}$$

The boundary $S_0 = 0$ has little importance as the traction due to the tip propagates over a finite characteristic length $L$ (see Section I E); for $\ell \gg L$ the base has no effect and can be considered frozen.

#### E. One-dimensional growth and tip-growth limit

We consider the simplified scenario of an axon towed longitudinally by a constant pulling force $\mathcal{F}$ [3]. Let $v(\sigma)$ denote the anterograde velocity field, with $\sigma := \ell - s$. At steady growth regime, $n$ and $v$ are functions of $\sigma$ only, with $n$ obeying $\mathrm{d}^2 n/\mathrm{d}\sigma^2 = n/L^2$ where $L := \sqrt{A\eta/\zeta}$. From the boundary condition $n(0) = \mathcal{F}$ we obtain:

$$n(\sigma) = \mathcal{F}\mathrm{e}^{-\sigma/L}, \tag{S9}$$

and

$$v(\sigma) = -\int \frac{n(\sigma)\,\mathrm{d}\sigma}{A\eta} = \mathcal{F}\mathrm{e}^{-\sigma/L}/\sqrt{A\eta\zeta}, \tag{S10}$$

which gives the tip velocity

$$V := v(0) = \mathcal{F}/\sqrt{A\eta\zeta}. \tag{S11}$$

### II. DUROTACTIC WRENCH

We compute the durotactic resultant and moment applied to a tract. We parameterize the cross section $\Omega$ by two coordinates $u$ (along the normal $\boldsymbol{\nu}$) and $z$ (in the out-of-plane direction). In general, $\mathcal{F}$ and $\mathcal{M}$ are given by:

$$\mathcal{F} = \vartheta \int_\Omega C(\boldsymbol{r} + u\boldsymbol{\nu})\,\mathrm{d}u\mathrm{d}z, \tag{S12}$$

$$\mathcal{M} = -\vartheta \int_\Omega C(\boldsymbol{r} + u\boldsymbol{\nu})\,u\,\mathrm{d}u\mathrm{d}z. \tag{S13}$$

#### A. Soft interface

In the case of a smooth substrate stiffness field, we can expand Eqs. (S12) and (S13) to leading order, to obtain

$$\mathcal{F} = A\vartheta C(\boldsymbol{r}) + \text{h.o.t.}, \tag{S14}$$

$$\mathcal{M} = -\vartheta C(\boldsymbol{r}) \int_\Omega u\mathrm{d}u\mathrm{d}z - I\vartheta\boldsymbol{\nu}\cdot\nabla C(\boldsymbol{r}) + \text{h.o.t.} \tag{S15}$$

The integral in the previous expression vanishes by taking the origin of $u$ and $z$ to be the centroid of the cross section:

$$\mathcal{M} = -I\vartheta\boldsymbol{\nu}\cdot\nabla C(\boldsymbol{r}) + \text{h.o.t.} \tag{S16}$$

#### B. Hard interface

In the case of a hard interface, the stiffness field is non-smooth, hence Eqs. (S14) and (S16) do not apply. The resultant and moment applied to a bundle sitting astride the interface depends on the proportion of the tip cross section that is located in each of the two zones. For a given configuration of the rod, we define $\chi(u) := 1$ if point $\boldsymbol{r} + u\boldsymbol{\nu}$ is in the $C_2$ zone, 0 otherwise. Then:

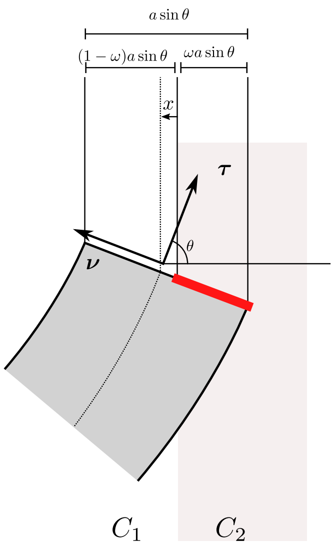


FIG. S1. Axon bundle straddling the interface, resulting in differential traction density.

$$C(\boldsymbol{r} + u\boldsymbol{\nu}) = C_1(1-\chi(u)) + C_2\chi(u). \tag{S17}$$

From Eqs. (S12) and (S13), we compute

$$\begin{aligned}\mathcal{F} &= \vartheta \int_\Omega (C_1(1-\chi(u)) + C_2\chi(u))\,\mathrm{d}u\mathrm{d}z \\ &= \vartheta C_1(A - J_1) + \vartheta C_2 J_1,\end{aligned} \tag{S18}$$

and

$$\begin{aligned}\mathcal{M} &= -\vartheta \int_\Omega (C_1(1-\chi(u)) + C_2\chi(u))\,u\,\mathrm{d}u\mathrm{d}z \\ &= -\vartheta\Delta C J_2,\end{aligned} \tag{S19}$$

where the two integrals $J_1 := \int_\Omega \chi(u)\,\mathrm{d}u\mathrm{d}z$ and $J_2 := \int_\Omega \chi(u)\,u\,\mathrm{d}u\mathrm{d}z$ depend on the shape of the cross section and the position and orientation of the tract. Assuming a rectangular cross section with width $a$ and thickness $b$, we calculate:

$$J_1 = \omega A, \tag{S20}$$

$$\begin{aligned}J_2 &= -\frac{a^2 b}{4}\,\mathrm{sgn}(\sin\theta) \int_{1-2\omega}^{1} h\,\mathrm{d}h \\ &= -\frac{6}{a} I\omega(1-\omega)\,\mathrm{sgn}(\sin\theta)\end{aligned} \tag{S21}$$

where $\omega = \int \chi/a \in [0,1]$ represents the proportion of the tip width that is inside the $C_2$ zone:

$$\omega(x,\theta) = \begin{cases} 0 & \text{if } x < a|\sin\theta|/2, \\ 1 & \text{if } x > a|\sin\theta|/2, \\ \frac{1}{2} + \frac{x}{a}|\csc\theta| & \text{otherwise.} \end{cases} \tag{S22}$$

From Eqs. (S18) to (S22), we obtain the exact expressions:

$$\mathcal{F} = A\vartheta\left(\left(1-\omega\right)C_1 + \omega C_2\right), \tag{S23}$$

$$\mathcal{M} = \frac{6}{a} I\vartheta\omega\left(1-\omega\right)\operatorname{sgn}\left(\sin\theta\right)\Delta C. \tag{S24}$$

### III. THEORETICAL ANALYSIS OF REFRACTION AND REFLECTION

In this section we provide a detailed theoretical analysis of the refraction/reflection problem (*Main text*, Eqs. (7) and (8)):

$$\frac{\mathrm{d}x}{\mathrm{d}s} = \cos\theta, \quad \frac{\mathrm{d}y}{\mathrm{d}s} = \sin\theta, \tag{S25}$$

$$\frac{\mathrm{d}\theta}{\mathrm{d}s} = \frac{3p}{2}\operatorname{sgn}\left(\sin\theta\right)\left(1-4x^2\csc^2\theta\right)_+, \tag{S26}$$

where $(x)_+ := \max(0, x)$ is the ramp function.

#### A. Symmetries

System (S25–S26) displays two interesting symmetries (see the phase portrait shown in Fig. 2B). First, invariance under $\theta \to -\theta$ translates mirror symmetry of the problem: each trajectory of the physical space possesses a twin that is obtained by reflection with respect to the interface normal. Second, invariance under $s \to -s$ and $\theta \to \pi + \theta$ signifies time-reversibility of the process, viz. a given path of the physical domain can be taken in both ways undifferentiably. Combining these two symmetries, we also see that tracts that do not make it across the interface – corresponding to those trajectories in the phase plane that make a U-turn – are reflected symmetrically with respect to the interface normal, i.e. $\theta_2 \equiv \pi - \theta_1$. Indeed, a reflection path crossing $\theta \equiv \pi/2$ is left unchanged (up to a change in direction) by translation $\theta \to \theta \pm \pi$ followed by reflection $\theta \to -\theta$ (Fig. 2B). This is only possible if the path is symmetric with respect to $\theta = \pi/2$, i.e. $\theta_2 \equiv \pi - \theta_1$.

#### B. Snell law

We derive the Snell law of bundle refraction (*Main text*, Eq. (10)). In virtue of the above-mentioned symmetries of System (7–8), we may restrict our study to $\theta \in [0, \pi]$. The influence region of the interface is the portion of the phase plane that is between the two $\theta$-nullclines $2x = \pm\sin\theta$. In this region, Eqs. (S25) and (S26) simplify to:

$$\frac{\mathrm{d}x}{\mathrm{d}s} = \cos\theta, \quad \frac{\mathrm{d}y}{\mathrm{d}s} = \sin\theta, \quad \frac{\mathrm{d}\theta}{\mathrm{d}s} = \frac{3p}{2}\left(1-4x^2\csc^2\theta\right). \tag{S27}$$

A first integral of this system is

$$\mathcal{I}(x,\theta) = \log\left(\sin\theta\right) + \sum_{i=1}^{3}\frac{4q_i^2-1}{12q_i^2-1}\log\left(x\csc\theta - q_i\right) \tag{S28}$$

where $\{q_1, q_2, q_3\}$ are the roots of the cubic $12pq^3 - 3pq + 2 = 0$ (Fig. S2A). A trajectory of Eq. (S27) intersects the interface boundaries at $(x,\theta) = (-\sin(\theta_1)/2, \theta_1)$ and $(x,\theta) = (\sin(\theta_2)/2, \theta_2)$. By definition, $\mathcal{I}$ is constant along the trajectories, which provides the Snell condition:

$$\mathcal{I}\left(-\sin\left(\theta_1\right)/2, \theta_1\right) = \mathcal{I}\left(\sin\left(\theta_2\right)/2, \theta_2\right). \tag{S29}$$

System (S27) has two fixed points $C := (-1/2, \pi/2)$ and $S := (1/2, \pi/2)$ (Fig. S2A). The point $C$ is a nonlinear center, which corresponds to reflection where the bundle grows tangent to the interface. The family of periodic trajectories around $C$ corresponds to the set of possible reflections. We see that $\theta_2 \equiv \pi - \theta_1$ satisfies Eq. (S29) for those solutions.

The other fixed point $S$ is a saddle point. Past this point, there are no periodic orbits and the system operates in a refraction regime. The critical orbit is the homoclinic orbit represented by a dashed line in the phase portrait (Fig. S2A). The incident angle $\theta^*$ associated with this orbit satisfies $\mathcal{I}(-\sin(\theta^*)/2, \theta^*) = \mathcal{I}(1/2, \pi/2)$, providing the critical angle $\theta^* = \arcsin\left(\exp\left(-c\left(p\right)\right)\right)$, with

$$c\left(p\right) := \sum_{i=1}^{3}\frac{4q_i^2-1}{12q_i^2-1}\log\left(\frac{2q_i+1}{2q_i-1}\right). \tag{S30}$$

The general Snell-like law of refraction is

$$\sin\theta_2/\sin\theta_1 = \mathrm{e}^{c(p)}. \tag{S31}$$

The $q_i$ are defined everywhere except at $p = 0$. However, it can be verified that $c(p) \to 0$ when $p \to 0$. In addition, $c'(p) \to 1$. Time-reversibility (Section III A) also implies that $c$ is an odd function, hence its Taylor polynomial only has odd non-zero terms. We deduce from the previous remarks $c(p) = p + \mathcal{O}\left(p^3\right)$ (Fig. S2B), hence the Snell approximation (*Main text*, Eq. (10)).

### IV. DURODUCT

The axon *duroduct* is modeled as a stiffness depletion $C(X,Y) = \bar{C} - \Delta C\exp\left(-Y^2/R^2\right)$. We derive the bundle curvature supplemented by a Gaussian noise $\epsilon$: $\kappa = -2py\cos\theta\exp\left(-y^2/R^2\right)/R^2 + \epsilon$. The travel time $\mathcal{T}$ associated with a path is estimated from the steady velocity $V$ (Eq. (S11)):

$$\mathcal{T} := \int \mathrm{d}t \approx \int \frac{\mathrm{d}s}{V\left(s\right)} = \bar{\mathcal{T}}\int\frac{\mathrm{d}s}{1-k\mathrm{e}^{-y^2/R^2}} \tag{S32}$$

where $\bar{\mathcal{T}} := \sqrt{\zeta\eta/A}/\bar{C}\vartheta$ is a characteristic time; $k := \Delta C/\bar{C}$ measures the intrinsic efficiency of the duroduct ($k = 0.25$ here).

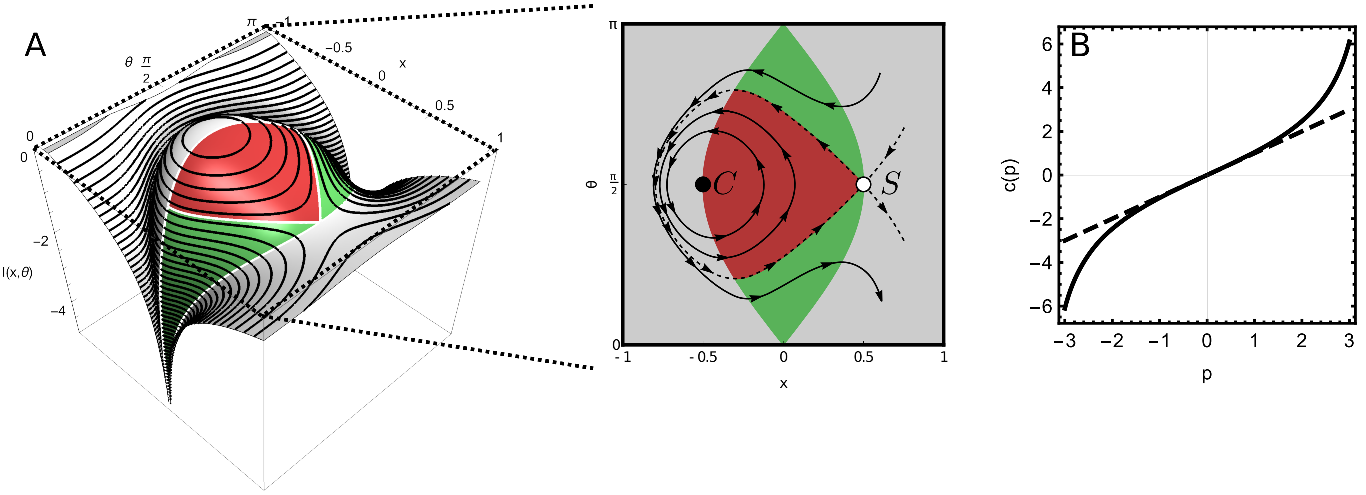


FIG. S2. (A) Plot of $\mathcal{I}(x,\theta)$ for $p = 0.5$ (Eq. (S28)) and phase portrait $x$-$\theta$ of System (S27). Light zone shows the implicit region $4x^2 \csc\theta > 1$ ($4x^2 \csc\theta = 1$ are the $\theta$-nullclines). Red and green domains depict the sets of reflection and refraction paths. (B) Plot of $c(p)$ vs. $p$ (Eq. (S30)). Dashed line shows leading-order approximation $c(p) \sim p$.

### V. XENOPUS OPTIC TRACT

#### A. Method

To simulate the growth of the xenopus optic tract studied in Refs. [10, 11], we construct a smooth stiffness field from measurements of the brain tissue stiffness obtained by *atomic force microscopy* [11], using a smooth surface interpolation (Fig. S3). The stiffness data consists of a $12 \times 7$ matrix, equivalent to a 240 µm×140 µm domain (pixel size: 20 µm). Since we here are only interested in torque effects, we normalize the substrate stiffness as

$$C^*(X,Y) = \frac{C(X,Y) - \min(C)}{\Delta C} \in [0,1] \qquad \text{(S33)}$$

with $\Delta C = \max(C) - \min(C) \approx 0.2$ kPa (the min and max are taken over the domain of interest). Eqs. (1) and (S16) simplify to $\kappa = -p\boldsymbol{\nu} \cdot \nabla C^*$, where $p = \vartheta \Delta C/E$ is the durotactic number for this problem.

Axons are initiated from the right or bottom edge of the domain of interest at a random distance from the bottom right corner chosen within a 20 µm interval. Similarly, initial angles are randomly chosen in $2\pi/3 \pm 10\%$.

#### B. Parameter estimates

To quantify the significance of the predicted durotactic turn, we derive a typical value for the durotactic number $p = \vartheta \Delta C/E$. The durotactic coupling coefficient $\vartheta$ is first estimated from single axon velocities measured *in vitro* by Koser *et al.* [10]. Two types of medium are considered here, with $C_{\text{soft}} = 0.1$ kPa and $C_{\text{stiff}} = 1$ kPa, resulting in two different velocities $V_{\text{stiff}} \approx 35$ µm.h$^{-1}$ and $V_{\text{soft}} \approx 22$ µm.h$^{-1}$. In our model, this velocity is proportional to the traction force (Eq. (S11)). The data suggests that

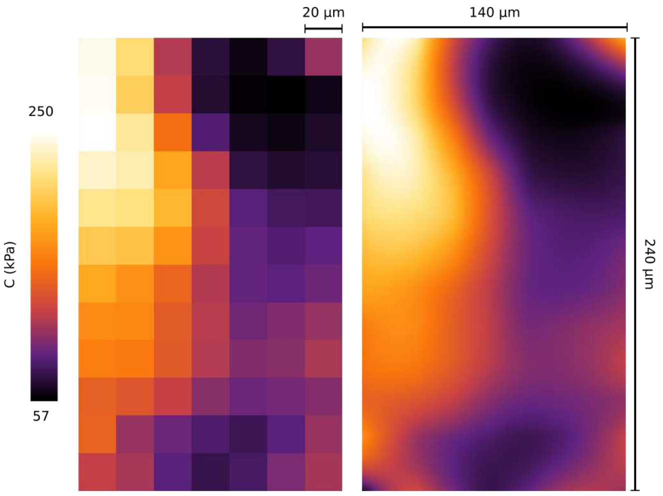


FIG. S3. Xenopus brain stiffness map. From left to right: discrete stiffness matrix obtained by atomic force microscopy [11], and smooth interpolation $C(X,Y)$.

the force velocity relationship has the general form $f = \vartheta C + f_0$ where $f_0$ is a constant. However, this constant does not contribute to the torque (Eq. (S13)) and can therefore be ignored for our analysis. We estimate the proportionality coefficient $\vartheta$ from

$$\vartheta = \frac{\sqrt{A\eta\zeta}\,(V_{\text{stiff}} - V_{\text{soft}})}{A\,(C_{\text{stiff}} - C_{\text{soft}})} = \frac{\eta}{L} \frac{V_{\text{stiff}} - V_{\text{soft}}}{C_{\text{stiff}} - C_{\text{soft}}}. \qquad \text{(S34)}$$

Using $L \approx 10$ µm for single axons (our own unpublished estimation) and $\eta = 1$–$10$ MPa.s [3], we find $\vartheta \approx 0.4$–$4$. A published measure for the Young's modulus is $E \approx 10$ kPa [6]. However, this corresponds to an extensional rigidity of single axons; the effective bending resistance of the bundle may be orders of magnitude lower due to the loose connections between axons, as indicated by the ability of bundles to form sharp bends [12]. Taking $E =$ 0.1–10 kPa and $\Delta C = 0.2$ kPa (Fig. S3), we find $p \approx$ 0.01–1.

### VI. IMPLEMENTATION

All simulations use standard builtin integration routines available in the software environment *Mathematica* 12 (`www.wolfram.com/mathematica/`). Codes will be made available upon acceptance for publication under license CC-BY-NC 4.0.

*


[1] D. E. Moulton, T. Lessinnes, and A. Goriely, Journal of the Mechanics and Physics of Solids **61**, 398 (2013).

[2] A. Goriely, *The mathematics and mechanics of biological growth* (Springer, 2017).
