## Supplementary figures and images for "An optic ray theory for durotactic axon guidance"

### Videos

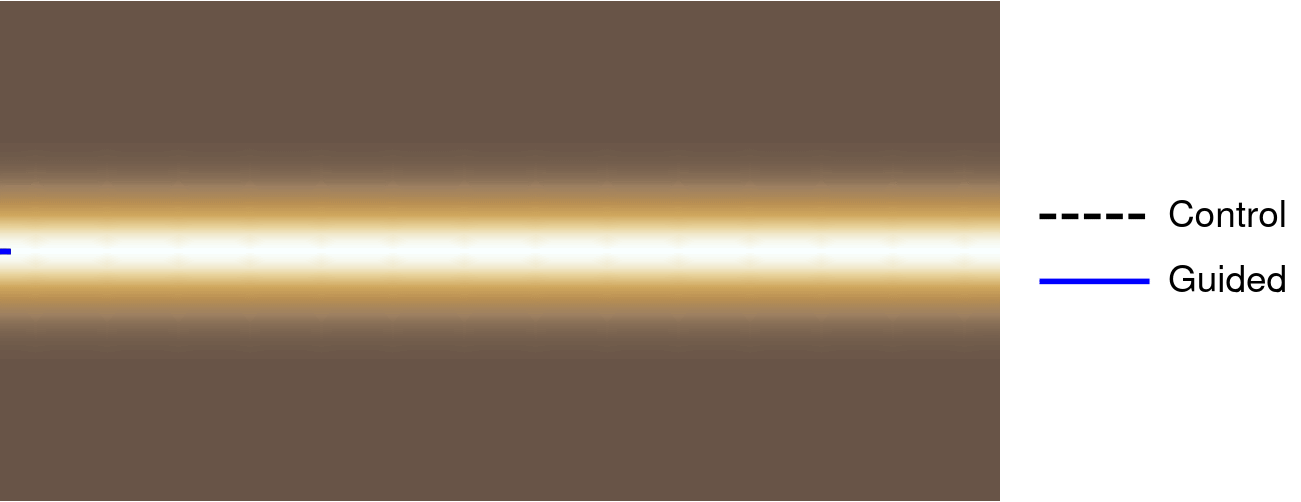
